# Ecological assembly of natural soundscapes in the Garajonay National Park (Spain)

**DOI:** 10.1101/2022.12.21.521439

**Authors:** Rüdiger Ortiz-Álvarez, Marta García-Puig, Leire Garate

**Affiliations:** Fundación Española para la Ciencia y la Tecnología (FECYT). 28100, Alcobendas, Madrid, Spain; Makara Divulga, G67543686, Travessera de les Corts 265, 08014, Barcelona, Spain; AZTI, Marine Research, Basque Research and Technology Alliance (BRTA), Herrera Kaia, Portualdea z/g, 20110 Pasaia, Gipuzkoa, Spain

## Abstract

Natural sound contains data about the ecology of animal populations, communities, and the full ecosystem, resulting from a complex evolution and varying according to the environment. Amongst the processes that are hypothesized to explain sound assemblages, or soundscapes, one is the acoustic niche hypothesis: sounds produced by species calling at the same time seek avoid overlapping, leading to an acoustic differentiation of signals. Soundscapes are more complex in the most pristine environments and show responses to habitat degradation and physical perturbations; hence here, we focus on La Gomera, in the Canary Islands (Spain). This island is the only location in Europe where primary cloud forests are well preserved and thrive on an island with varied orography, microclimates, disturbances, and vegetation types. In this article, we adapted a method to quantify the importance of acoustic niche partitioning and also the opposite process: acoustic aggregation. To do so, we explored soundscapes at different temporal scales in forests with variable degrees of perturbation and maturity. A secondary goal of this report is to compare how soundscapes could differ in an area affected by a wildfire, and undisturbed equivalents, in summer in winter, seasons with contrasting temperatures and wind regimes. We conclude that tracking faunal activity and behavior through soundscape monitoring could be a piece of useful complementary information to guide conservation decisions and future restoration efforts in the Garajonay National Park (La Gomera).

**Highlights:**

○ The mature forest and the transitional forest are similar in terms of sound levels, frequencies, and dynamics, although the transitional forest had higher sound levels of the lower frequencies (higher NDSI).
○ The mature forest has stronger and more active assembly mechanisms (i.e., acoustic aggregation and acoustic niche partitioning) and a higher acoustic diversity index (ADI) than the transitional forest. Assembly organizes differently in both locations.
○ In both the mature and the transitional forests, we found an inverse relationship between the ADI and acoustic niche partitioning. At the highest diversities (more frequency bands occupied), the weakest is the temporal avoidance of calls with similar frequencies.
○ The vulnerability of the burned location is highest during the harsh summer, but in winter, it hosts a community very similar to the intact vegetation in terms of ADI and sound levels.

## Introduction

Sound is an emergent property inherent to any ecosystem. The combination of animal signals, geological features, and anthropogenic impacts non-randomly arrange to form soundscapes ^1^, which are the focus study of two study fields: soundscape ecology ^2^ and its umbrella discipline ecoacoustics ^3^. One of the main goals of these fields is the understanding of the underlying processes behind natural sound organization. Biological signals arrange with maximum complexity in undisturbed ecosystems ^4,5^, fruit of a complex and multifactorial evolution ^6–8^, and subject to a changing environment ^9,10^. Under the lens of community ecology, ecological assembly is a central question ^11^. However, dynamics and rules are still being understood at the soundscape level.

Amongst the potential drivers of the soundscape assembly, the acoustic niche partitioning hypothesis has often been examined ^1,3,12^. Under this hypothesis, species arrange their calling activity to avoid frequency overlap by calling at different frequencies or separating their activity into daily and seasonal activity patterns, or by occupying differently the physical space ^12^. Although there is significant evidence agreeing with this hypothesis ^3,13^, there are contradicting examples where calling activity seems to concentrate and arrange around a particular frequency instead ^14^. In addition, a study showed an effect of relaxed competition on song evolution on islands compared to the mainland ^5^, remarking the importance of quantifying the importance of the acoustic niche hypothesis and other phenomena behind soundscape dynamics.

This article aims at unveiling the ecological assembly behind the natural soundscapes of Garajonay National Park in La Gomera (Canary Islands, Spain). This island is the only spot in Europe where mature cloud laurel forests -*Laurisilva*- have been preserved since the Pliocene/Pleistocene transition 2-3 million years ago ^15^, and the only location in the Canary Islands with extant primary forests ^16^. Furthermore, the varied orography of La Gomera island is home to heterogeneous plant community assemblages organized through distinct microclimates. Such a situation makes La Gomera a suitable location to explore the factors behind soundscape assembly within very short distances. In particular, to explore how the activity of sonoriferous communities arrange through habitats with different degrees of degradation and during different seasons.

In La Gomera, vegetation degradation is associated with a number of long-term conservation challenges. Some are location-specific, and others are the fruit of a global context. Globally, there is a 25-year trend toward less diverse soundscapes ^9^. Climate change has been highlighted as a stressor of natural soundscapes because alterations in temperature and humidity can compromise species survival, behavior, and sound propagation ^10^. Also, there is an increasing risk of wildfires, which are a major threat to La Gomera forests ^17^. Indeed, in 2012 on the southern slope of the island there was a big forest fire burning 3617 ha, including ~750 ha of protected vegetation ^18^, mostly *fayal-brezal* (transitional vegetation to Laurisilva in drier conditions). Ever since, authorities’ efforts have promoted the natural restoration of the area and implemented restoration activities. Whilst there are strategies to characterize the successional trajectories of Laurel forests and associated vegetations ^19^, there is still a need to understand the behavior and colonization of sonoriferous fauna during the recovery ^17^.

We aim at contributing with a new layer of ecoacoustic information to the complex ecosystem of Garajonay National Park. We explored soundscapes by covering different temporal scales (hours, days, seasons) in different habitats with variable degrees of perturbation and maturity. Particularly, we have examined in detail acoustic dynamics during the spring of 2020 in two Laurisilva forest spots with different levels of maturity: a fully mature forest, and a *fayal-brezal* in transition to Laurisilva. In these protected locations we are particularly focusing on understanding seasonal succession, quantifying acoustic niche partitioning, and inferring other possible assembly rules. A secondary goal of this article is to compare how soundscapes could differ in an area affected by a wildfire, and undisturbed equivalents, in summer in winter, seasons with contrasting temperatures and wind regimes. We show how acoustic monitoring to study biodiversity and habitat health is an emerging approach that is fast, cost-effective, non-invasive, and that provides information about the management efforts led by the National Park authorities. In addition to the conservation perspective, we highlight the ecoacoustics research potential of La Gomera to add to the big picture rules of natural soundscape assemblages.

## Methods

### Habitats, seasons and recording scheme

We focus on two different major goals. Under the first goal, to examine in depth the acoustic assembly, we compare two Laurisilva forests in the northern slope, with different levels of maturity (Meriga and El Cedro), and their seasonal transition from Spring towards Summer (**Table 1**). The arrangement of plant species and their maximum canopy heights was different, with the mature forest dominated by taller *Persea indica* trees, and in the other hand, the less mature forest was mixed with *Laurus azorica* and *Myrica faya* among others. For the second goal, we compare a burned location (Cabezo de Mocanillo) with equivalent intact counterparts in the southern slope (El Bailadero and Las Creces) in two seasons: summer and winter (**Table 1**). In both cases we used autonomous SM4 recorders (Wildlife Acoustics), set to record at 48000 Hz, 4 minutes every hour, and 30 minutes adjusted automatically every day around dusk and dawn choruses.

**Table 1.** Study locations and their associated vegetations, for two different goals: (A) Laurisilva with different levels of maturity. (B) Comparing a burned location with an intact equivalent.

| Location | Vegetation | Slope | Coordinates | Season | Start date | Days |
| --- | --- | --- | --- | --- | --- | --- |
| Meriga Forest (A) | Mature laurisilva | Northern | 28°08'36.3"N 17°14'19.6"W | Spring | May 11th 2020 | 53 |
| El Cedro Forest (A) | Transition to Laurisilva | Northern | 28°07'45.7"N 17°13'19.3"W | Spring | May 11th 2020 | 51 |
| Bailadero (B) | Fayal brezal intact (near ravine) | Southern | 28°07'22.1"N 17°12'28.7"W | Summer | Aug 15th 2020 | 13 |
| Cabezo M. (B) | Fayal brezal burned | Southern | 28°07'34.7"N 17°16'16.8"W | Summer | Aug 15th 2020 | 33 |
| Las Creces (B) | Fayal brezal intact | Southern | 28°08'01.1"N 17°17'24.8"W | Winter | Dec 27th 2020 | 6 |
| Cabezo M. (B) | Fayal brezal burned | Southern | 28°07'34.7"N 17°16'16.8"W | Winter | Dec 27th 2020 | 7 |

### Spectral profiles

Spectral composition was calculated by first, filtering each channel for each sample removing the signal below 5%, then the spectrum for each channel was calculated using the *spec* function in Seewave package ^20^, using a window length of 512, FFT function for faster calculation, returning the Probability Mass Function (PMF). The spectrums of left and right channels were averaged with a resolution of two decimals (kHz) to obtain a single spectrum per time and location.

### Sound levels and bioacoustics indexes

Noise levels for each fragment were quantified in Kaleidoscope PRO v. 5.4.2, adjusting gain levels according to the automatic process of SM4 recorders, and given with a reference of 0 dB = 20 microPascals. Sound level averages were estimated by transforming dB to a linear scale. For the calculation of bioacoustics indexes we also used the implementation within Kaleidoscope PRO, focusing on the Acoustic Diversity Index (ADI) and Normalized Difference Soundscape Index (NDSI). ADI ^21^ was selected to quantify occupied frequency niches: it divides the frequency spectrum into 10 bins (500 Hz each) and takes the proportion of the signals in each bin above a −50dBFS threshold. Then, ADI is the result of the Shannon index applied to these bins. NDSI ^22^ is the ratio between common anthropophony and geophony frequencies versus biophony frequencies. In this article, we used a 20-1000 Hz range for the geophony and anthropophony, and a 1000-10000 Hz range for the biophony, given that anthropophony was not intense, particularly because part of the recordings happened during the COVID-19 restrictions ^23,24^.

### Temporal associations of call clusters and temporal frequency overlap

#### Adapted method

To test the acoustic niche hypothesis in the northern slope locations, Meriga and El Cedro, we adapted a strategy often used in ecology to test the phylogenetic relatedness of a species trait ^25^. That strategy measures the relatedness between the value of a species trait of interest, and how phylogenetically close are a group of species. Here, instead of species, we use clusters derived from the automatic clustering analysis of the the Kaleidoscope PRO v. 5.4.2 software. Instead of phylogenies, we use phylograms based on cluster co-occurrences and co-exclusions. And the trait of interest to test an association with is ‘call mean frequency’. By this framework, we could approximate how calls with similar frequencies tend to aggregate or to avoidance, and how both phenomena could be acting simultaneously.

#### Constructing a cluster/time table

First, we required a table of signals assigned to different clusters. We conducted the analysis under default parameters, detecting signals between 250 to 12000 Hz, and obtaining a table of putative calls assigned to different clusters. These raw clusters can be considered groups of similar calls, likely coming from a single species. However, this method retrieves a high number of false positives ^26^, and specific individual assignments can be wary without manual inspection. We use these automatic clusters as proxies of the acoustic community, but not as 100% verified species-specific calls, by quantifying the number of vocalizations of the three top matches, weighting them by the inverse distance of assignment to each cluster (*Cluster ‘abundance’ = ∑_for each cluster and time_ Max distance – call to cluster distance*).

#### Estimate call associations

With a table of cluster ‘abundances’ for each location (Meriga vs. El Cedro) divided into 30-second time fragments, we calculated the positive and negative associations of these clusters by using a network-based approach as implemented in the wTO package ^27^ in R. This method allows to keep negative correlations ^28^ and to remove random links (Spearman correlations) by block-bootstrapping ^27^ (250 bootstraps and a determined lag of 42 in the *acf* function). *P values* were adjusted with the Benjamini & Hochberg correction, keeping only associations with *p (adjusted)* < 0.001.

#### Input files to estimate the K statistic

The resulting weighted associations (absolute values) were transformed into lists of dissimilarity matrices, using the inverted weights as dissimilarities. The dissimilarity matrices were the basis to construct WPGMA trees ^29^ as implemented in *hclust*. Trees constructed using positive associations have co-occurring clusters in the nearest branches, while trees constructed using negative associations have co-excluding clusters in the nearest branches. The goal was to evaluate if there is a relationship between the phylograms, where branch lengths represent the weights of co-occurrences and co-exclusions -i.e., two close branches are call clusters that tend to cooccur or that avoid each other-) and ‘call cluster mean frequency’. Through the R package Picante ^25^, we calculated the K statistic ^30^, a Brownian motion-based metric of the strength of the relatedness signal. The statistical significance of K is estimated by permuting trait values across the tips of a tree. A significant signal here indicates the tendency for calling clusters with similar frequencies towards aggregation or partitioning.

Code is available from the corresponding author by request.

### Multivariate statistics and ordinations: general assembly rules

To examine the relationships between acoustic properties in the multivariate acoustic space, we performed a PCA analysis by using means at the daily level. Acoustic variables were (1) Sound levels of 20-1000 Hz, characterized as geophony, (2) sound levels of 2000-8000 Hz characterized as biophony, bioacoustics indexes (ADI and NDSI), K statistics (based in co-occurrences, and co-exclusions, separately), and the axis resulting from an nMDS analysis, that represent the variation in spectral composition, based on Bray-Curtis dissimilarities between the spectrums. In addition, three environmental/metadata variables were incorporated: temperature (as measured internally within the SM4 recorders), sun altitude (representing a seasonal variation), and moon light, to consider potential variations during nights with more or less light, and also seasonal variations, estimated with R package *suncalc* ^31^.

## Results and Discussion

### The acoustic space during late Spring in Laurisilva forests

We are first comparing a mature forest thriving in a valley (Meriga Forest) with a secondary forest in a hillside, in transition to mature *Laurisilva* (El Cedro). Average sound levels had similar intensity (dB) and dynamics in both locations, every hour and every day from May to July (**Figure 1**). Still, these levels were slightly higher in the secondary forest, possibly indicating higher exposure to winds, or noise from one of the island’s main roads which normally have lower frequencies (see boxplots in **Figure 4A** for levels by frequency range). Hourly, both locations had dawn and dusk choruses, standard peaks of activity associated with sun position ^32^, and an additional smaller peak at 15:00 (**Figure 1C)**.

**Figure 1.**
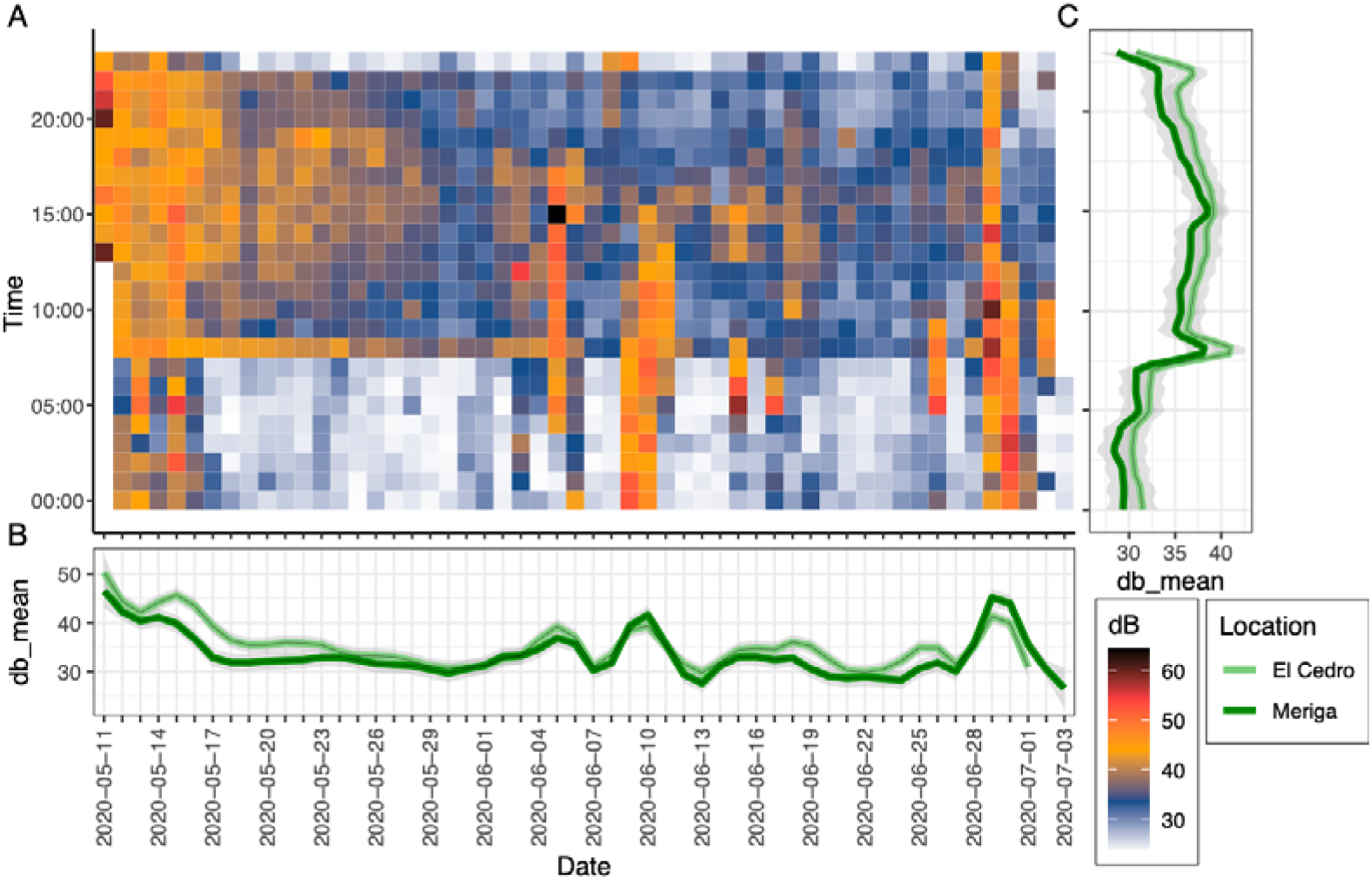
(**A**) Average sound pressure levels (dB) in the Meriga Forest and El Cedro Forest, for the frequencies between 125 Hz and 12699 Hz, (**B**) Seasonal variation of the sound pressure levels, (**C**) daily variation of the sound pressure levels.

In both locations, during the time series, sound levels were higher during May (**Figure 1B**). Despite three particular meteorological events increasing sound levels and altering frequency compositions (**Figure 2**), there were consistent changes. Until the end of May, there was a soundscape dominated by calls at ~2.1kHz, likely dominated by the blackbird *Turdus merula*, an abundant bird in many habitats in La Gomera ^33^, changing towards calls at ~5kHz, and decreasing their activity towards a more heterogeneous acoustic space by the start of the summer, with some activity at ~7.5kHz.

**Figure 2.**
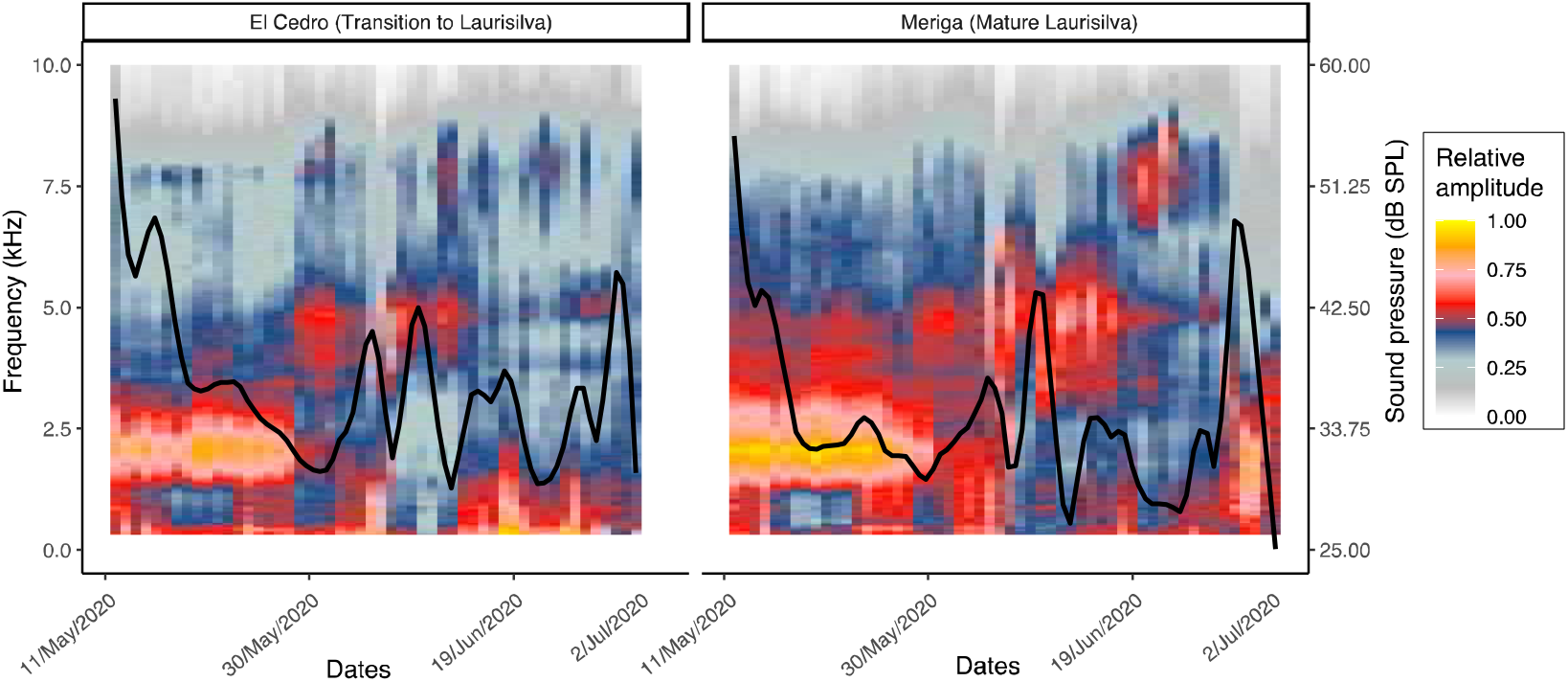
Seasonal spectrogram of frequencies between 20 Hz to 10 kHz. The secondary y-axis and the black line show the sound pressure (dB) variation.

### Acoustic aggregation vs. partitioning of calls with similar frequencies

To quantify the impact of the acoustic niche hypothesis we propose an automatized approach that can be used to compare different habitats, and both possibilities: frequency aggregation and frequency partitioning. We use the *K* statistic ^30^, a metric that quantifies the relatedness signal between calls and their frequencies (i.e., the higher the value, the higher the association between call frequency and temporal overlap/avoidance).

#### Call aggregation

In both forests, calls with similar frequencies tend to occur together. Such a result comes from the observation that although both the temporal aggregation (based on co-occurrences), and avoidance (based on co-exclusions) were significantly associated with mean frequencies (**Figure 3**), the K statistic had higher values when calculated using co-occurrences. Furthermore, when examining the K statistic to compare aggregation in both habitats at different time scales, we observe marked differences. In particular, estimating the K statistic with all the species of each location together at the same time, call aggregation was stronger in the transitory vegetation (El Cedro) than in the mature Laurisilva (Meriga). Conversely, examining in detail the temporal dynamics of the K statistic and call aggregation, aggregation was stronger in the mature Laurisilva (Meriga), at the beginning of the time-series when Blackbirds dominate the soundscape (**Figure 3B**), and between dawn and dusk (**Figure 3C**). Such dominance of call aggregation might be mediating interspecific competition among species with similar ecological niches, as previously observed in the Amazon rainforest ^14^, or simply a synchronized vocal behavior of multiple species ^34^. Or perhaps, we are observing the convergence of calls from only a few species, given the low avian diversity of La Gomera compared to tropical rainforests or mainlands ^33^, so we might be observing clashes between individuals and/or populations.

**Figure 3.**
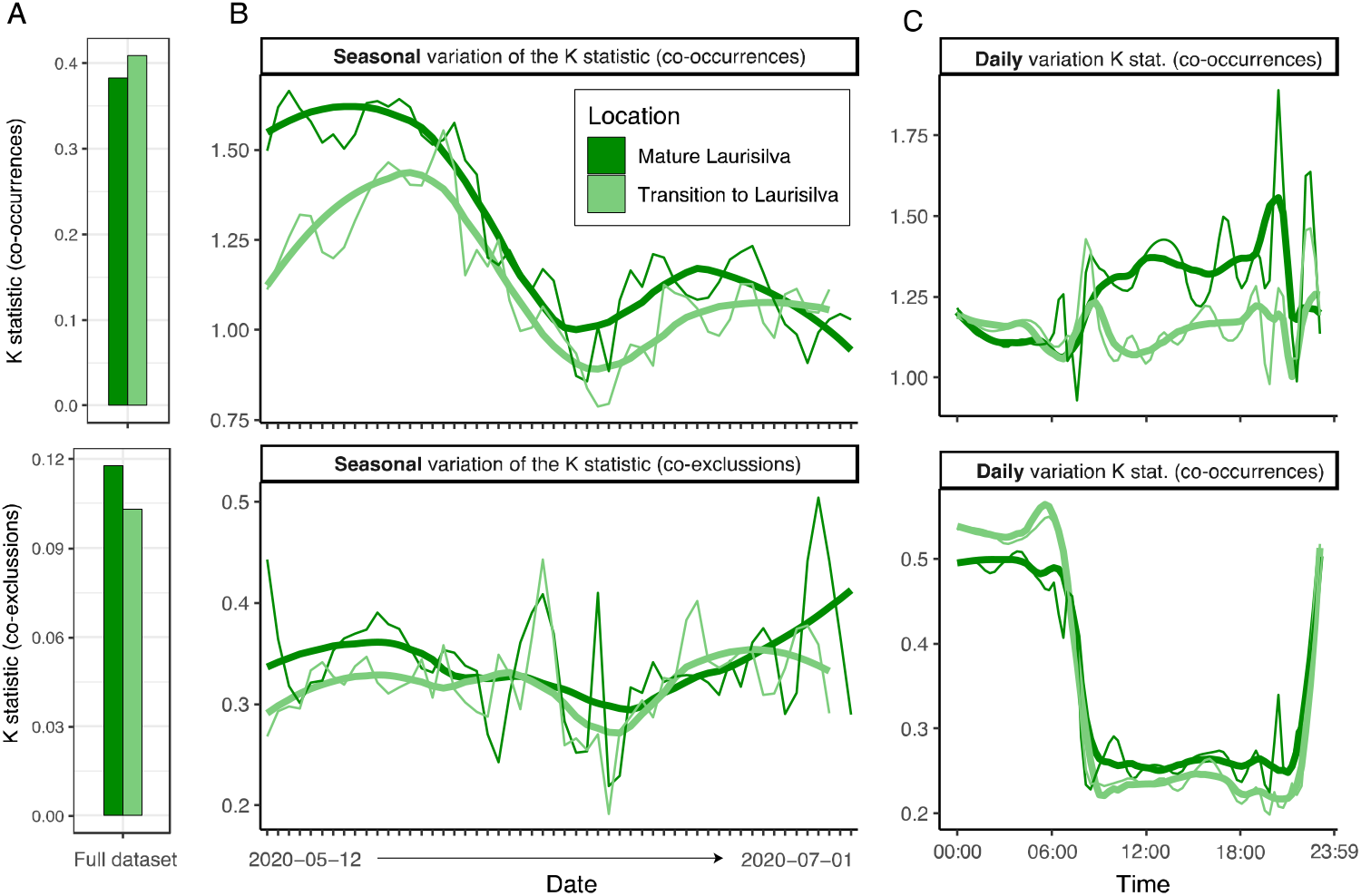
Quantification of the acoustic niche partitioning and aggregation. (**A**) Full dataset, (**B**) Seasonal variation, (**C**) Daily variation.

#### Call avoidance

Calls with similar frequencies avoid each other, with lower K statistics but still significantly, indicating a role of acoustic niche partitioning (ANP). The lower impact of ANP might be due to a reduced competition of species on islands, since islands may impose fewer constraints on sonoriferous fauna ^5^. Such partitioning was slightly higher in the mature Laurisilva (**Figure 3A and 3B**), the oldest vegetation, than in the younger habitat, agreeing with the overall expectation ^3,13^. Indeed, higher ANP is associated with increased species richness and increased levels of acoustic interference ^5^ and spectral partitioning ^35^. By inspecting the temporal dynamics of acoustic niche partitioning, we observe that it is particularly clear during the daytime, with avoidance peaks coinciding with dawn and dusk choruses (**Figure 3C)**.

#### Simultaneous processes

Studies have highlighted cases where acoustic partitioning is detected, and others have not ^3,34^. Here we point out that both phenomena might coexist in the island, showing different seasonal patterns. Our results are a good hint that co-occurrence/co-exclusion analyses transformed into trees, combined with a test of a trait phylogenetic signal (K statistic), could be a fruitful way to quantify the acoustic niche hypothesis and acoustic aggregation at different spatial and temporal scales.

### Habitat differences on the acoustic assemblages

Acoustic metrics can detect even minor differences between habitats, since physical changes are likely to be mirrored in the local soundscape ^36–38^. Here, taking all the acoustic metrics together: sound levels, frequency composition, our approximation to quantify differences on acoustic aggregation/partitioning, bioacoustics indexes and environmental metadata, we aimed to discern part of the processes behind soundscape assembly in two closely related habitats (**Figure 4**).

**Figure 4.**
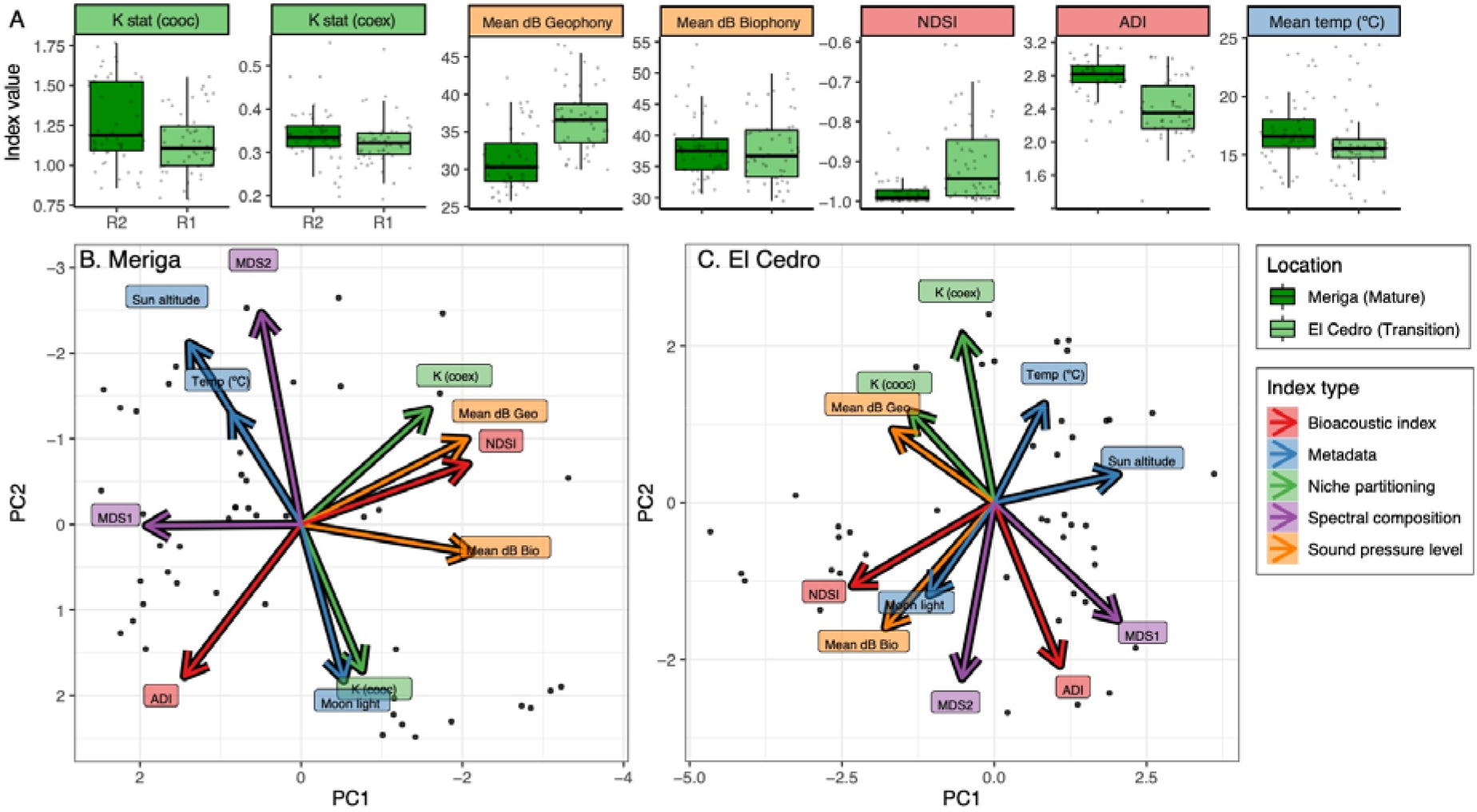
(**A**) Boxplots showing differences between the locations in terms of K statistics, mean sound levels, bioacoustics indexes, and temperature. Data were combined into two Principal Component Analyses (PCA) for the (**B**) Meriga Forest and (**C**) El Cedro Forest.

#### ADI vs. temporal partitioning

We observe an inverse relationship between acoustic diversity (ADI) and acoustic niche partitioning (ANP) independently in both locations. This result means that at the highest acoustic diversities, with more frequency bands occupied, the weakest was the avoidance of calls with similar frequencies at the same time. It is often considered that partitioning is increased when most species are found in a community by temporal over-dispersal or adjusting their frequencies ^35,39^. The inverse relationship we observe in each location separately (**Figure 4B and 4C**) might be showing a process on a different scale: when diverse calls arrange in the frequency spectrum, there might be no need for temporal separation.

#### Spectrum vs. temporal aggregation

Call aggregation has a strong relationship with the spectral composition (MDS2 in Meriga, and MDS1 in El Cedro), indicating that frequency spectral composition might depend on the ecological processes that lead to temporal aggregation of calls with similar frequencies ^14^, such as vocal synchronization ^34^. Perhaps, depending on the activity of the avian community or its seasonal turnover.

#### Habitat differences

Despite the similarities of both vegetations, the ecology of communication is not identical (**Figure 4B and 4C**). ADI is substantially higher in the mature forest, Meriga, despite the similar dB levels of biophony in both ecosystems. Another notable example is an inverse relationship such dB levels and the recorded temperature in El Cedro, while being unrelated in Meriga. Or the increased NDSI in El Cedro, perhaps due to increased wind exposure or due to being more exposed to one of the island’s main roads. We hint at the mosaic of environmental situations that might shape differently the island’s soundscapes, differentiating even very similar ecosystems. Standardized and grid sampling designs could shed more light into what are the environmental factors behind soundscape differences ^40^.

### Wildfire of the southern slope

Here we compared sound levels (**Figure 5**) and acoustic diversity (**Figure 6**) of a burned location vs. an intact location in two seasons: summer and winter. In general, we observed similar temporal dynamics with varying sound levels. During summer, the burned area was much quieter than its intact equivalent and did not show any dawn or dusk choruses, while during winter, both areas were very similar and displayed the dawn and dusk choruses similarly (**Figure 5A**). In summer, a quiet but present dawn chorus occurred only in the intact vegetation. Such result suggest that the increased heat of the summer highlighted the vulnerabilities of the burned area, possibly impeding shelter of sonoriferous species due to the lack of canopy since canopy structure can drive differences in faunal behavior ^41,42^. Despite this vulnerability, at night, populations of crickets and cicadas, singing at high frequencies (**Figure 5A)**, matched the dynamics of the intact area, highlighting the presence of active insect populations. We do not know how the environmental variables affect insects, although there are examples showing that increased wind exposure ^43^ and temperature ^44^ can modify insect calling activity.

**Figure 5.**
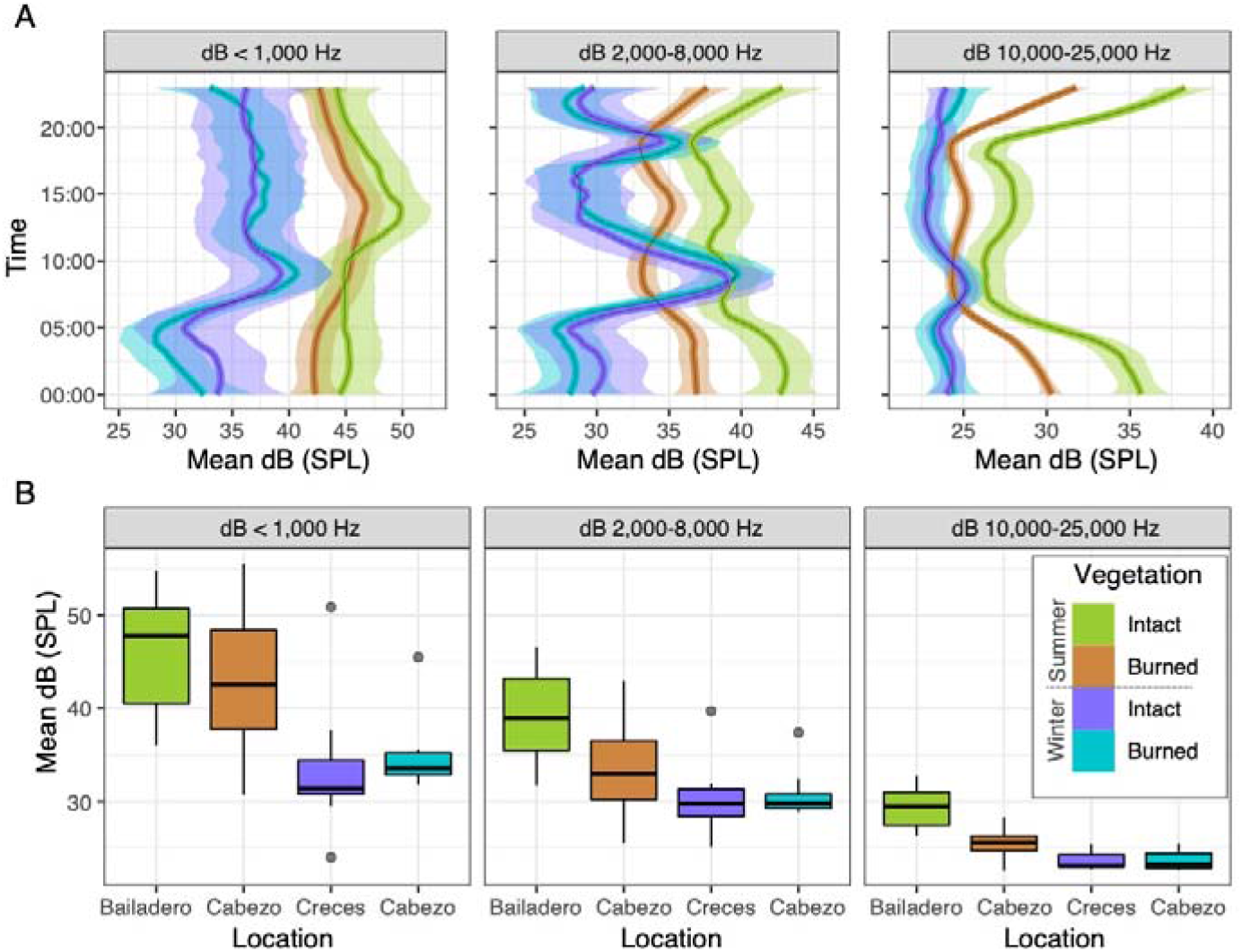
(**A**) Daily variation of sound pressure levels, and (B) Mean dB sound pressure levels at three frequency ranges: <1000 Hz, between 2000-8000 Hz and between 10000-25000 Hz. Sound levels are shown comparing a burned area vs. an intact area, during summer and during winter.

**Figure 6.**
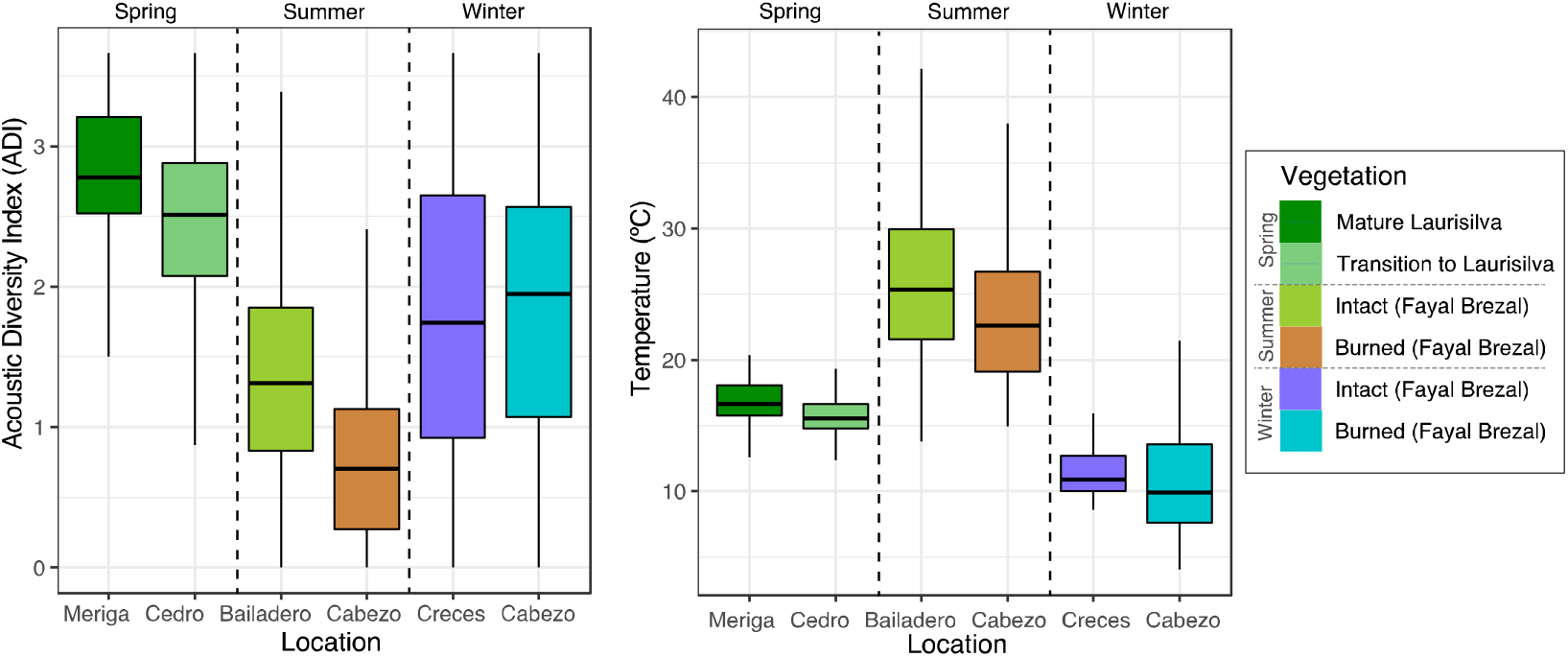
(**A**) Acoustic Diversity Index (ADI), and (**B**) temperature by type of vegetation and season (Spring, Summer and Winter).

Regarding acoustic diversity, based on a single index (ADI), in winter both locations showed similar values (**Figure 6**). Conversely, in summer, ADI was decreased in the burned location. Indeed, the burned location in summer had the lowest diversity of all the seasons and locations explored in this article. Lower acoustic diversities in burned areas and particularly during summer, have been observed in other locations undergoing natural recovery ^45^. Furthermore, the physical attributes of habitats can force the evolution and adaptation of animal calls to maximize their propagation ^46,47^, so perhaps soundscapes will vary during the long-term recovery of the area according to vegetation changes. In general, burned areas in laurel forests of the Canary Islands have the potential to recover, although it is a long process and intermediate vegetation stages seem to favor fire propagation ^17^.

## Awknowledgements

This work has been funded by the National Geographic Society (grant NGS-64645R-19). Fieldwork permits were granted by the authorities of Garajonay National Park. The authors thank Luis L. Pinto, Dr. Cèlia Sitjà and Park authorities: Ángel B. Fernández and Antonio Zamorano Benavides for fieldwork assistance and logistics. We thank Dr. Laurel B. Symes for her support to grant and for her helpful comments during the process. The authors declare no conflict of interest.

